# An updated assembly strategy helps parsing the cryptic mitochondrial genome evolution in plants

**DOI:** 10.1101/2021.03.02.433321

**Authors:** Yanlei Feng, Xiaoguo Xiang, Zhixi Fu, Ronghui Pan, Xiaohua Jin

## Abstract

Although plant mitogenomes are small in size, their variations are no less than any other complex genomes. They are under rapid structure and size changes. These characters make the assembly a great challenge. This caused two intertwined problems, a slow growth of known mitogenomes and a poor knowledge of their evolution. In many species, mitogenome becomes the last genome that undeciphered. To have a better understanding of these two questions, we developed a strategy using short sequencing reads and combining current tools and manual steps to get high quality mitogenomes. This strategy allowed us to assembled 23 complete mitogenomes from 5 families in Fagales. Our large-scale comparative genomic analyses indicated the composition of mitogenomes is very mosaic that “horizontal transfers” can be from almost all taxa in seed plants. The largest mitogenome contains more homologous DNA with other Fagales, rather than unique sequences. Besides of real HGTs, sometimes mitovirus, nuclear insertions and other third-part DNA could also produce HGT-like sequences, accounting partially for the unusual evolutionary trajectories, including the cryptic size expansion in *Carpinus*. Mitochondrial plasmid was also found. Its lower GC content indicates that it may be only an interphase of a foreign DNA before accepting by the main chromosome. Our methods and results provide new insights into the assembly and mechanisms of mitogenome evolution.

## Introduction

Organelle genomes are always known as small size and conserved structure, such as the mitochondrial genome (mitogenome) in animals and plastid genome (plastome). Somehow, mitogenome in angiosperms (or seed plants) is totally different. Its size is largely expanded but also varies significantly among species, with no less than 200 Kb and up to 11 Mb (Sloan et al., 2012; exception see Skippington et al., 2015). Alone with the expansion, duplications, plastid-derived insertions (referred as mitochondrial plastid insertions, MTPTs) and horizontal DNA transfers (HGTs) are common. DNA double strand breaks are rampant and structure is extremely dynamic (Davila et al., 2011; Christensen, 2018), chromosome in many species becomes linear, multipartite, branched or a mixture of them (Alverson et al., 2011a; Cheng et al., 2017; Kozik et al., 2019). Even close relatives or individuals of the same species may have different mitogenomes.

These special characters hinder the complete and high-quality assembly of mitogenome. Using published relative as a reference is a general strategy of assembling plastomes. However, the poor conservation and uncertain chromosome structure of mitogenome make this way impossible. On the other hand, *de novo* assembly is not easy, either. The dispersed repeats always make the contigs very fragmental. What’s more, the coverage of plastome is normally much higher than mitogenome. Therefore, how to get the precise content of MTPTs remains challenging. So far, there are softwares for plastome and animal mitogenome, such as GetOrganelle, Novoplasty and mitoZ (Dierckxsens et al., 2017; Meng et al., 2019; Jin et al., 2020). Some scientists used bacterial genome tools instead, like Unicycle (Wick et al., 2017). But none of their performance on plant mitogenome is satisfactory. The tools specially plant mitogenome is still blank.

A direct consequence of this difficulty is the slow growth of this area. In angiosperms, the sequenced mitogenomes are much less than plastomes and even nuclear genomes (a brief count in species level, mitogenomes: no more than 300, NCBI; plastome: more than 4000, NCBI; nuclear genome: more than 500, https://www.plabipd.de; till 2021 January). In many species, mitogenome becomes the last genome that not been deciphered. In addition, the publications are mostly about one or several species. Large-scale comparisons of mitogenome are still rare, remaining many mysteries about its evolution. Recently, many researchers start to use PacBio or Nanopore long reads. It’s better than short Next-Generation Sequencing (NGS) reads. But likewise, long repeats and long MTPTs couldn’t be solved perfectly as well. Its high price also limits the number of the sampling. The past years our working on plastomes and nuclear genomes has accumulated a mess of NGS data. If there is a way can reuse the data to get high-quality mitogenomes, it will save us a lot of money and time.

Fagales are an order belonging to Rosids of eudicots. Based on the APG system, they contain more than 1,000 species in 7 families and 33 genera (Sennikov et al., 2016). Many of them are very important for the ecology as well as food supply. They include many best-know trees, *e.g.*, beaches, oaks and birches, and many of them can offer nuts and fruits, *e.g.*, walnuts, chestnuts, hazels and bayberries. Fagales are also a diverse and ecologically dominant plant group that have nitrogen-fixing species. So far, at least 24 genomes were sequenced (https://www.plabipd.de) and 150 plastomes were released in Fagales, while only three mitogenomes were known. One is *Betula pendula* (GenBank ID: LT855379, Betulaceae, without annotation) from the WGS project, but in the publication no real content about it (Salojärvi et al., 2017); another is *Quercus variabilis* (GenBank ID: MN199236, Fagaceae) with only simple descriptions (Bi et al., 2019); the third is *Fagus sylvatica* (GenBank ID: MT446430, Fagaceae) published recently (Mader et al., 2020). Parsing their mitogenomes, the last unknown genetic material, is crucial for insight into their adaptive capacity, genetic and genomic resources.

In this study, we summarized the difficulties in mitogenome assembly with short reads and explored a strategy to overcome them. We used raw short reads from NCBI SRA (https://www.ncbi.nlm.nih.gov/sra) to assemble mitogenomes of 23 species, including 16 genera from 5 families of Fagales, covering almost half of the total genera and 71% of the total families respectively. By these high quality complete mitogenomes, we analyzed their differences by comparative genomics, especially size variation. This is one of the largest sampling in mitogenome studies heretofore.

## Results

### An updated strategy to get complete mitogenomes

Most projects sequenced the full DNA instead of isolating mitochondrion. Currently there is no way to get the mitochondrial reads directly from the total. We used *de novo* assembly, then found mitochondrial contigs by genes and coverage (details see Methods). Mitogenomes of angiosperms normally don’t contain AT-rich regions, the read coverage is quite balanced without gap (from known mitogenomes and experience). During assembling, the extension stops when meets repeats and MTPTs, i.e., contigs usually end either by repeats (repetitive ends) or MTPTs (MTPT ends). Firstly, we solved all the repeat ends by coverage (Figure 1, the coverage of a two-copy repeat pair is doubled). It may introduce artificial rearrangements between repeat pairs longer than the insert length. Afterwards, we mapped all MTPT ends to its plastome, and found the most potential connections. At last, we used the reads to correct MTPTs (Figure S1) and assembly (details see Methods).

**Figure 1:**
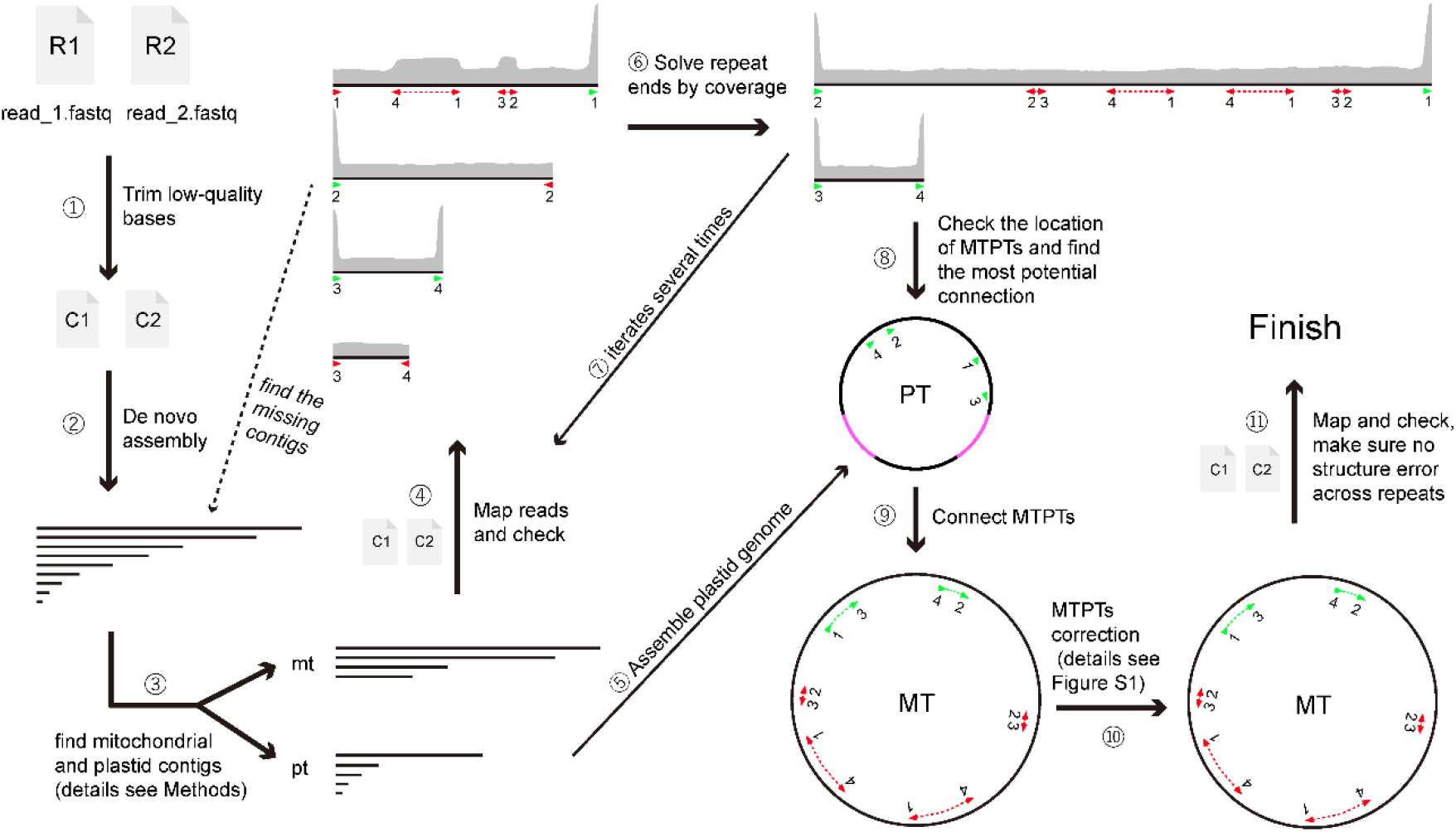
Workflow of our assembly strategy. Red and green arrows indicate repetitive and MTPT ends, respectively. The direction of the arrow points to the direction the end should extend to. Same number under red or green arrows means repeat or MTPT pair. R1, R2: raw reads; C1, C2: clean reads.

### Mitogenome assembly, structure and completeness assessment

The coverage of the assembled mitogenomes ranges from 33 to 174 (Table S1). 13 of the 23 species got circle(s), while the other 10 got one or multiple linear chromosomes (Table 1). In the circular assemblies, if there is a pair of repeats longer than the insert length between chromosomes, we merged the chromosomes into one through the repeats. The multi-circle mitogenome means no mergeable long repeat shared by chromosomes. In principle, if we connect all the repetitive and plastid ends, we can get circular mitogenome(s). However, sometimes the repetitive end pointed to an unstable, gradually decreasing coverage (Figure S2), we could not find out how to connect them; or the MTPT end failed to find the other pair (like *Corylus* chromosome 2). Other potential problems hinder we got circular chromosomes including: (1) In *Alnus* the repeats are rampant, many repeats duplicate three or more times. That’s the main reason why it’s so fragmented; (2) The coverage of *Bet. platyphylla* and *Lithocarpus* is quite unstable (it’s likely lower around AT-rich region). We don’t know if that was affected by a different sequencing machine. The reason why not circular and potential errors are summarized in Table S2. We believe that circular chromosomes are closer to an “ideal” complete mitogenome. Those non-circle assemblies should have no big problem about the DNA content, but repeat and MTPT length might be affected (Table S2).

**Table 1.** Basic information of Fagales mitogenomes. In column “Chr”, the number means chromosome number, while “C” and “L” behind represent “circular” and “linear”, respectively. The two species in red were assemblies from other projects.

| Species | Family | length (bp) | Chr | GC (%) | CDS | tRNA | rRNA | Repeat (bp) | MTPT (bp) |
| --- | --- | --- | --- | --- | --- | --- | --- | --- | --- |
| *Alnus glutinosa* | Betulaceae | 629389 | 16L | 45.44 | 35 | 18 | 3 | 8581 | 29856 |
| *Betula pendula* | Betulaceae | 581505 | 1L | 45.52 | 36 | 19 | 3 | 3703 | 31908 |
| *Betula platyphylla* | Betulaceae | 581519 | 2L | 45.53 | 36 | 19 | 3 | 3724 | 32032 |
| *Carpinus cordata* | Betulaceae | 922154 | 3C | 44.97 | 34 | 20 | 3 | 16557 | 46637 |
| *Corylus avellana* | Betulaceae | 635030 | 2L | 44.58 | 35 | 22 | 3 | 39128 | 60416 |
| *Ostrya chinensis* | Betulaceae | 688786 | 1L | 45.23 | 34 | 18 | 3 | 2601 | 31075 |
| *Ostryopsis nobilis* | Betulaceae | 669332 | 1C | 45.21 | 35 | 18 | 3 | 4810 | 24710 |
| *Casuarina equisetifolia* | Casuarinaceae | 492230 | 2C | 44.15 | 35 | 23 | 3 | 2431 | 66400 |
| *Casuarina glauca* | Casuarinaceae | 445851 | 2C | 44.92 | 34 | 19 | 3 | 1758 | 21776 |
| *Fagus sylvatica* | Fagaceae | 504715 | 1C | 45.85 | 34 | 17 | 3 | 2702 | 4615 |
| *Castanea mollissima* | Fagaceae | 388038 | 1C | 45.67 | 36 | 20 | 3 | 10888 | 10169 |
| *Lithocarpus fenestratus* | Fagaceae | 485396 | 6L | 45.76 | 35 | 17 | 3 | 13254 | 7122 |
| *Quercus robur* | Fagaceae | 390878 | 1C | 45.92 | 35 | 17 | 3 | 22006 | 8140 |
| *Quercus suber* | Fagaceae | 478989 | 1L | 45.85 | 36 | 19 | 3 | 26635 | 4967 |
| *Quercus variabilis* | Fagaceae | 412886 | 1C | 45.76 | 36 | 17 | 3 | 20747 | 4537 |
| *Cyclocarya paliurus* | Juglandaceae | 628759 | 1C | 44.89 | 37 | 19 | 3 | 2422 | 34143 |
| *Juglans cathayensis* | Juglandaceae | 740307 | 1C | 45.16 | 37 | 20 | 3 | 25019 | 22632 |
| *Juglans hindsii* | Juglandaceae | 716397 | 1C | 45.25 | 36 | 17 | 3 | 2118 | 12514 |
| *Juglans microcarpa* | Juglandaceae | 623287 | 1L | 45.2 | 35 | 18 | 3 | 6984 | 12448 |
| *Juglans nigra* | Juglandaceae | 716680 | 1C | 45.26 | 37 | 17 | 3 | 2022 | 12667 |
| *Juglans regia* | Juglandaceae | 775914 | 3L | 45.19 | 35 | 17 | 3 | 16232 | 15813 |
| *Juglans sigillata* | Juglandaceae | 778034 | 1L | 44.87 | 36 | 17 | 3 | 19731 | 34425 |
| *Platycarya strobilacea* | Juglandaceae | 502903 | 1C | 45.26 | 37 | 19 | 3 | 3253 | 28548 |
| *Pterocarya stenoptera* | Juglandaceae | 603233 | 1L | 45.35 | 36 | 17 | 3 | 5269 | 3524 |
| *Morella rubra* | Myricaceae | 523452 | 2C | 45.33 | 38 | 18 | 3 | 5668 | 9312 |

During our preparation of this project, the mitogenome of *Fagus sylvatica* was published (Mader et al., 2020). By using both long and short reads, the mitogenome was assembled into a single circle of 504,715 bp in length. Our assembly of *Fagus sylvatica* got the same size, with nearly 100% identical content - only two bases are different (difference between individuals rather than mis-assembly). The only disparity of the two version is a “inversion” between a 900-bp repeat. The insert length of our *Fagus* data is 450 bp (Table S1). This kind of disparity is expected. The work of Mader et al. (2020) has proved the practicability and reliability of our methods. These two independent projects at last got nearly the same mitogenome, it also proved, at least in some species, the mitogenome is still well preserved among individuals.

### Mitogenome size and content

The basic characters of our assemblies, as well as *Bet. pendula* and *Que. variabilis*, were listed in Table 1. Mitogenomes size in Casuarinaceae, Fagaceae and Myricaceae still resemble their distant relatives from Rosales or Fabales (400 and 480 Kb in average, respectively, based on the current data on NCBI). While in Betulaceae and Juglandaceae the size is obviously expanded. *Car. cordata* from Betulaceae has the largest size (922 Kb), around 2.4 times longer than the shortest, chestnut *Castanea* from Fugalaceae (388 Kb). However, other members in Betulaceae don’t have comparable length. Within each family, DNA content doesn’t differ much, though the structure is highly rearranged (Figure S3). The mitogenomes among families are less similar that many sequences have no homologs in others (Figure 3).

The proportion of repeats in Fagales mitogenomes is small, normally less than 3% and no more than 6.2% of the total length (Table 1). In Betulaceae, short repeats less than 200 bp obvious more, especially in *Alnus* (Table S3). Interestingly, some are highly copied, just like the “Bpu” sequences found in *Cycas* (Chaw et al., 2008). Situation is same to MTPTs. Only two species have MTPTs more than 6%, *Cas. equisetifolia* (13.5%) and *Corylus* (9.5%, maybe not exact, see Table S2).

The gene content of Fagales resembles other angiosperms. The 24 “core” protein coding genes (*atp1*, *4*, *6*, *8* and *9*, *ccmB*, *C*, *Fc* and *Fn*, *cob*, *cox1*-*3*, *nad1*-7, *9* and *4L*, *matR* and *mttB*), three ribosomal RNA genes (*rrn5*, *rrnS* and *rrnL*) and two succinate dehydrogenase subunit genes (*sdh3* and *sdh4*) are well preserved. Like many plants, the conservation of ribosomal protein genes is poor. Only 5 of them, *rpl5*, *rpl10*, *rps1*, *rps4* and *rps12*, exist in all. While the rests show different degrees of losses in different species (Figure 2). The *rps11* of *Betula* and *Corylus* was reported from horizontal gene transfer (HGT; Bergthorsson et al., 2003). We confirmed the conclusion, meanwhile expended the scope. Five of the seven Betulaceae species have this gene, with identity around 100%, either having open reading frame (ORF) or not. Blasting against NCBI nr database displayed it is more similar to monocots or the basal core angiosperms, *e.g.*, *Triantha glutinosa* (KX808303, Alismatales) and *Liriodendron tulipifera* (NC_021152, Magnoliales), which is also consistent with the former results (Bergthorsson et al., 2003). That indicate the HGT of *rps11* may happened to the common ancestor of Betulaceae, followed with differential losses in some species (Figure 2). The other gene, *rps13*, also only presents in minority. But BLAST result showed it similar to many Rosales, which may not support it’s an HGT.

**Figure 2.**
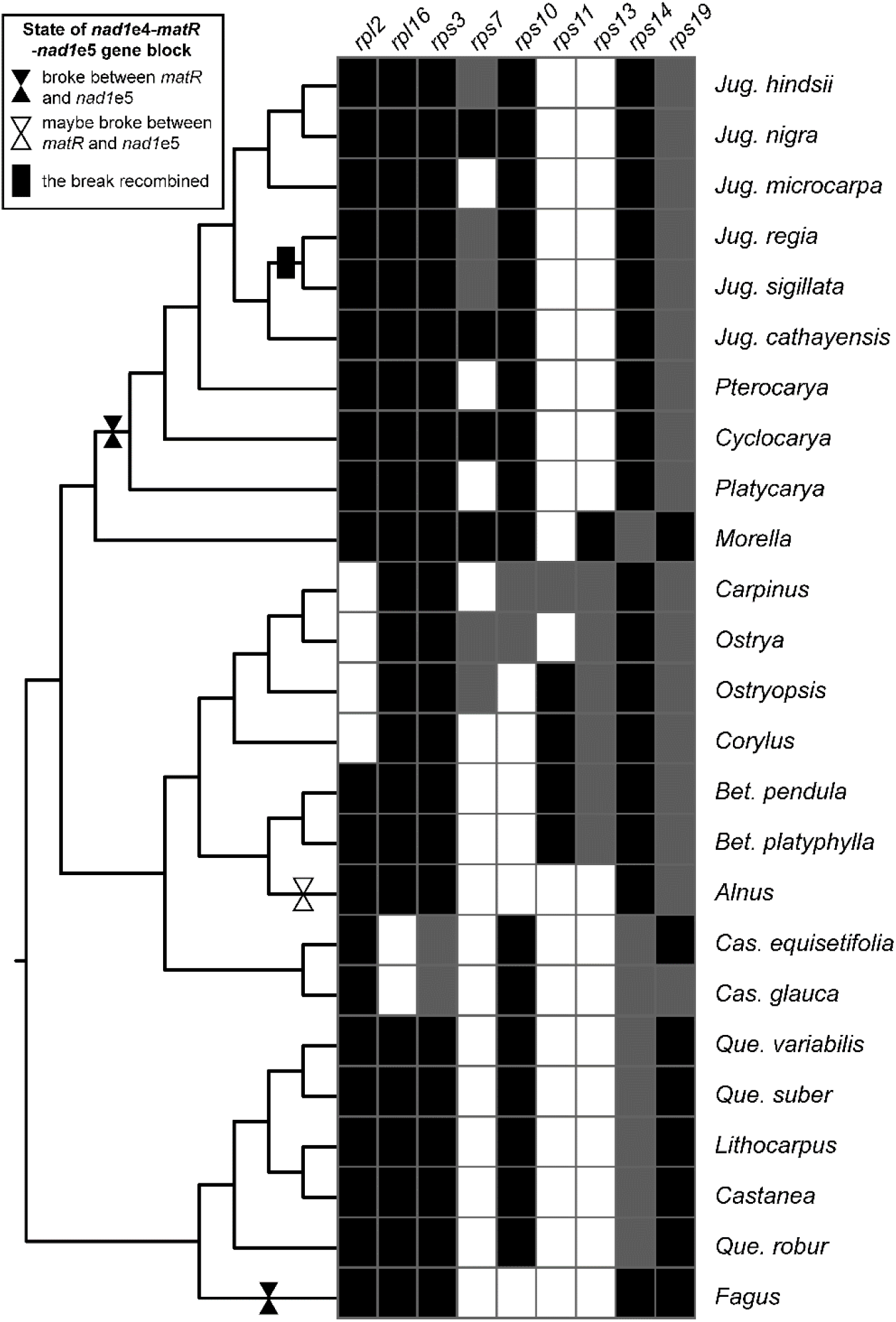
Genes that involved in variation. Black, grey and blank in the grids indicate a gene is intact, pseudo and missing, respectively. The breaks and reunion of *nad1*e4-*matR*-*nad1*e5 block is marked on the tree branches.

**Figure 3.**
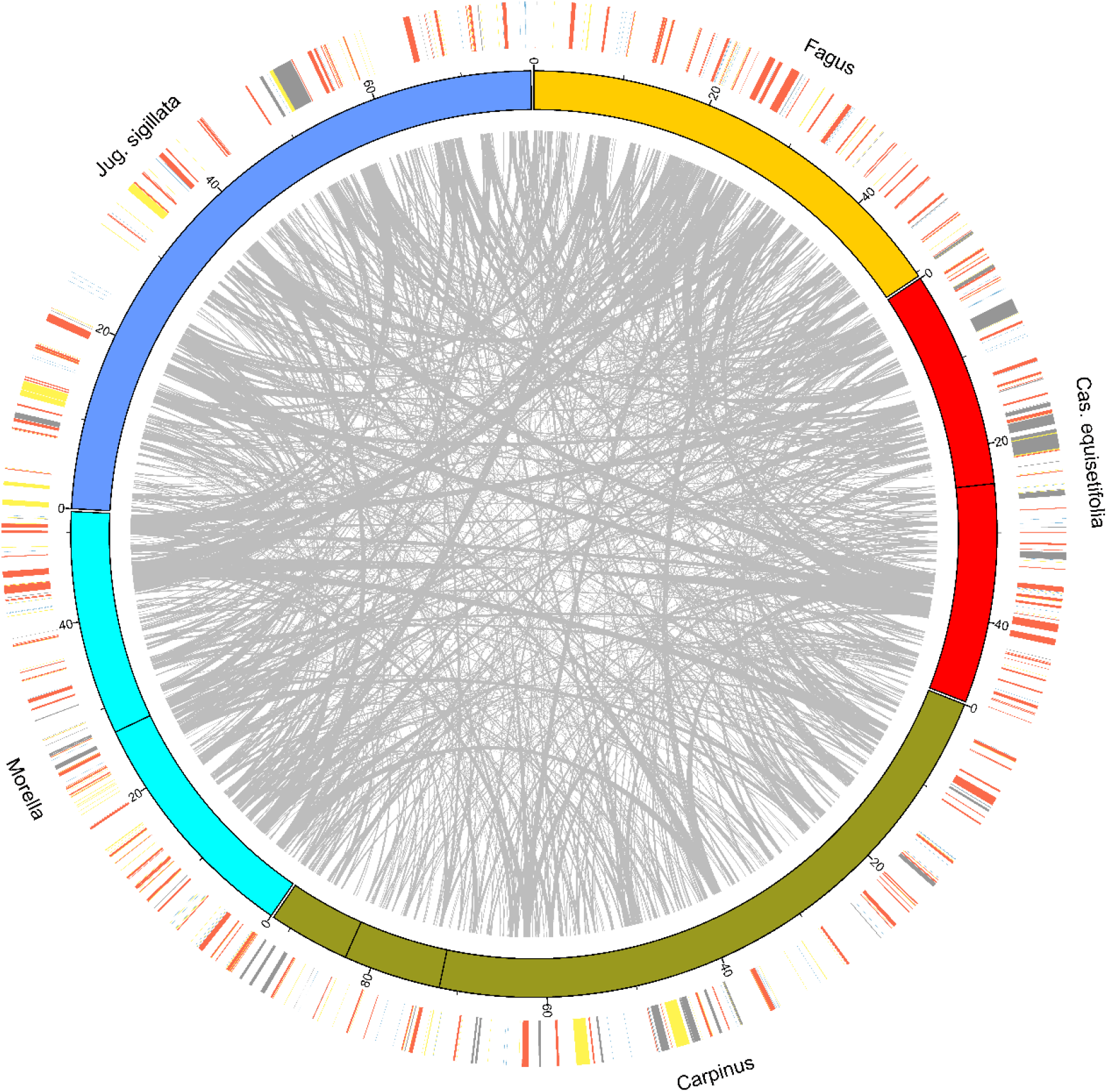
Circos plot of species from five families. The outer ring shows the position of protein genes and rRNA (red), tRNA (blue), repeats (yellow) and MTPTs (grey).

The *nad1* exon 4 (*nad1*e4), *matR* and *nad1*e5 form a gene colinear block in most angiosperms. The block broke between *matR* and *nad1*e5 at least two times in Fagales (Figure 2). One happened to *Fagus*, another happened to Juglandaceae (Figure 2). Surprisingly, in *Jug. sigillata* and *Jug. Regia*, this break was recovered. In Juglandaceae, a repeat around 110 bp is located within *matR* and *nad1*e5. Rearrangement between the repeat pairs may flip-flop that triggering the break and recovery. Whereas no repeat can be found in the corresponding area of *Fagus*. Its break maybe happened through nonhomologous recombination (Davila et al., 2011; Christensen, 2013). *Alnus* may also have this break. Since our assembly of *Alnus* is fragmental, further conclusion needs to be verified.

### Mitochondrial plasmid

In *Carpinus*, a very small circular mitochondrial plasmid was found, with only 2,888 bp in length. It has similar coverage (or slightly lower) to the main mitogenome. Its GC content is 37.6%, much lower than normal mtDNA (Table 1). It shares no homologs with other mitogenomes in angiosperms, except a *ca.* 240 bp plastid-like region. It can be fully covered by *Corylus avellana* or *Carpinus fangiana* nuclear sequences from different chromosomes. The plasmid has two large ORFs, ORF244 (732 bp) and ORF162 (486 bp). BLASTP against nr database indicate homologs of ORF244 were annotated as putative F-box protein in many species in angiosperms, including a nearly full-length matching in *Arabidopsis* (AT1G74875, identical 34%). homologs of ORF162 were annotated as “factor of DNA methylation 4” in many Rosids. Although we are not sure if the ORFs are expressed, evidences seem enough to prove the plasmid is from nucleus. The plasmid was not included in the total length or analyses of *Carpinus*.

### Phylogeny

The mitochondrial and plastid matrices are 31,551 and 69,243 bp long with 750 and 6,495 parsimony-informative characters, respectively. The phylogeny of Fagales based on chloroplast data (Figure S4b) showed that most of phylogenetic relationships are consistent with previous studies, but the relationships among Juglandaceae was quite different with Mu et al. (2020). While the phylogeny of Fagales based on mitochondrial tree (Figure S4a) showed that positions of the families in the plastid tree consists with previous work (Li et al., 2004). But the position of Myricaceae is incongruent with Xiang et al. (2014) and Xing et al. (2014). *Quercus* was reported as polyphyly in plastid trees but as monophyly in nuclear genes. Hybridization, introgression or incomplete lineage sorting possibly happened (Manos et al., 2008; Simeone et al., 2016). Both mitochondrial and plastid tree in our results showed a polyphyletic relationship of *Quercus*, which is also agreed with previous studies. The mitochondrial tree is poorly supported. It is almost expected since only few signals can be detected in the matrix. In phylogeny reconstruction, mitochondrial genes may be not effective enough in order or lower level. Hence, in this study we used the plastid tree for exhibition and discussion.

### Genus-specific DNA and mosaic origin

The detectable repeats and MTPTs are not sufficient to explain the size variation (Table 1). To further explore what caused the divergence, we isolated the specific DNA of each genus, *i.e.*, sequences have no homologs in other genus in Fagales. *Que. robur* was not merged into *Quercus*. To our surprise, the longest *Car. cordata* does not contain strikingly long specific sequences (Table S3); while *Casuarina*, whose size is relatively small, has the most (Table S3). Plant mitogenomes prone to absorbing foreign DNA. To analyze if these genus-specific sequences were from other species, we searched them against NCBI nr database. Only best-hits were counted and then incorporate into orders (Figure 4; Table S3-S4). Except those had no BLAST results, results related to most lineages in seed plants and most from mitogenomes (Figure 4).

**Figure 4.**
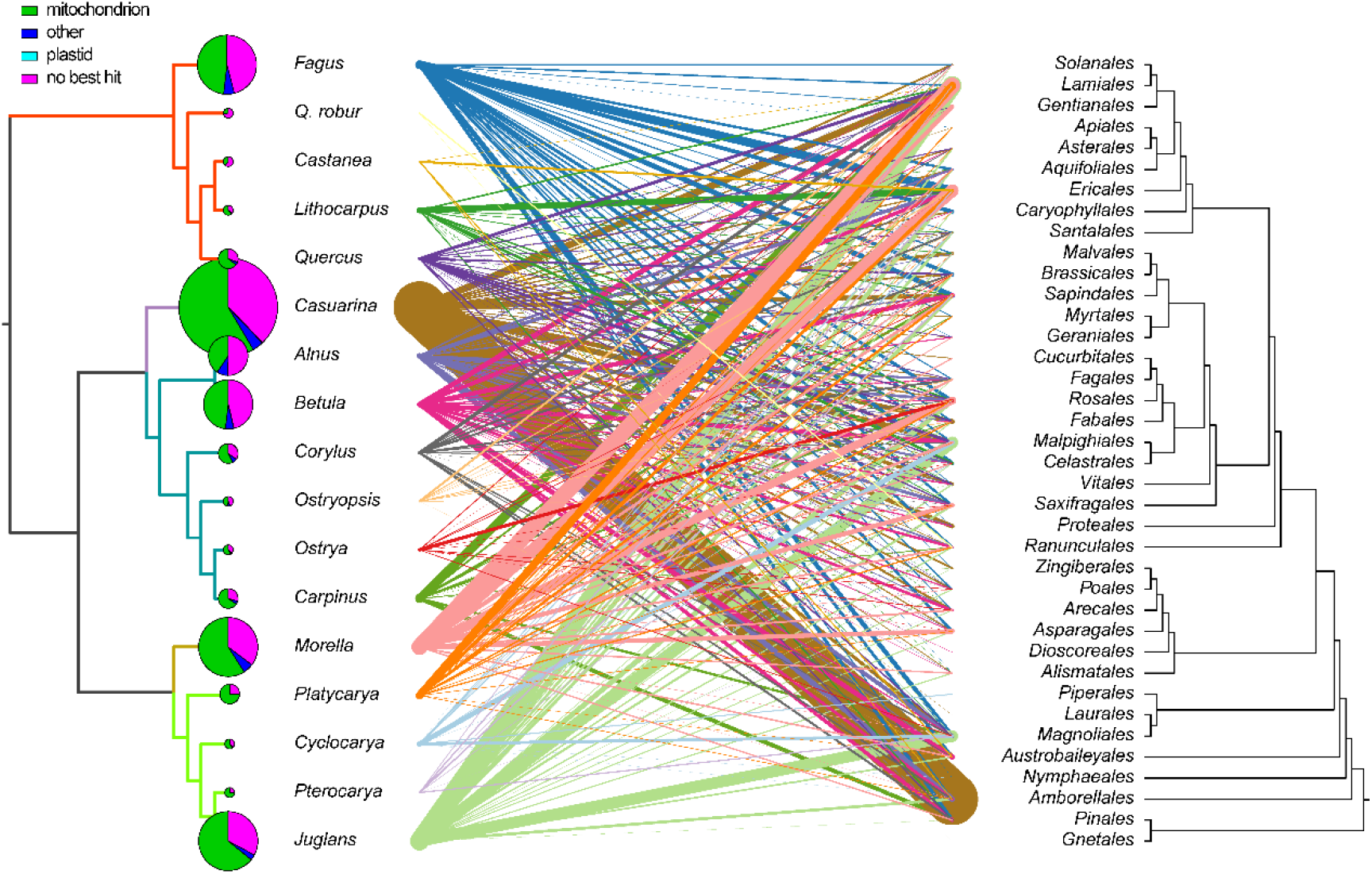
Components of the best-hits of genus-specific DNA. Best-hits of genus-specific DNA between species in Fagales and hit taxa (combined into orders) were connected by lines. Line thickness indicates total length and each species used the same color. Pie charts on the left show the proportions of mitochondrial, plastid and other hits. Pie size represents the total genus-specific DNA length. This plot was based on Table S4 and S5.

Mitovirus-like sequences were found in many species of Fagales, including a nearly full length in two *Betula*, and *ca.* 1500 bp in *Castanea* (Figure 5). Mitovirus (belongs to family Narnaviridae) is a kind of positive single-strand RNA virus that replicates in host mitochondria. Its genome is only 2.1 – 4.4 Kb in length and contains a single ORF of a viral RNA-dependent RNA polymerase presumably (RdRP) required for replication (Nibert, 2017). The phylogeny of Fagales mitovirus-like sequences is incongruent with the species tree (Figure 5), which indicates these mitovirus-like sequences were not integrated into Fagales by one single event.

**Figure 5.**
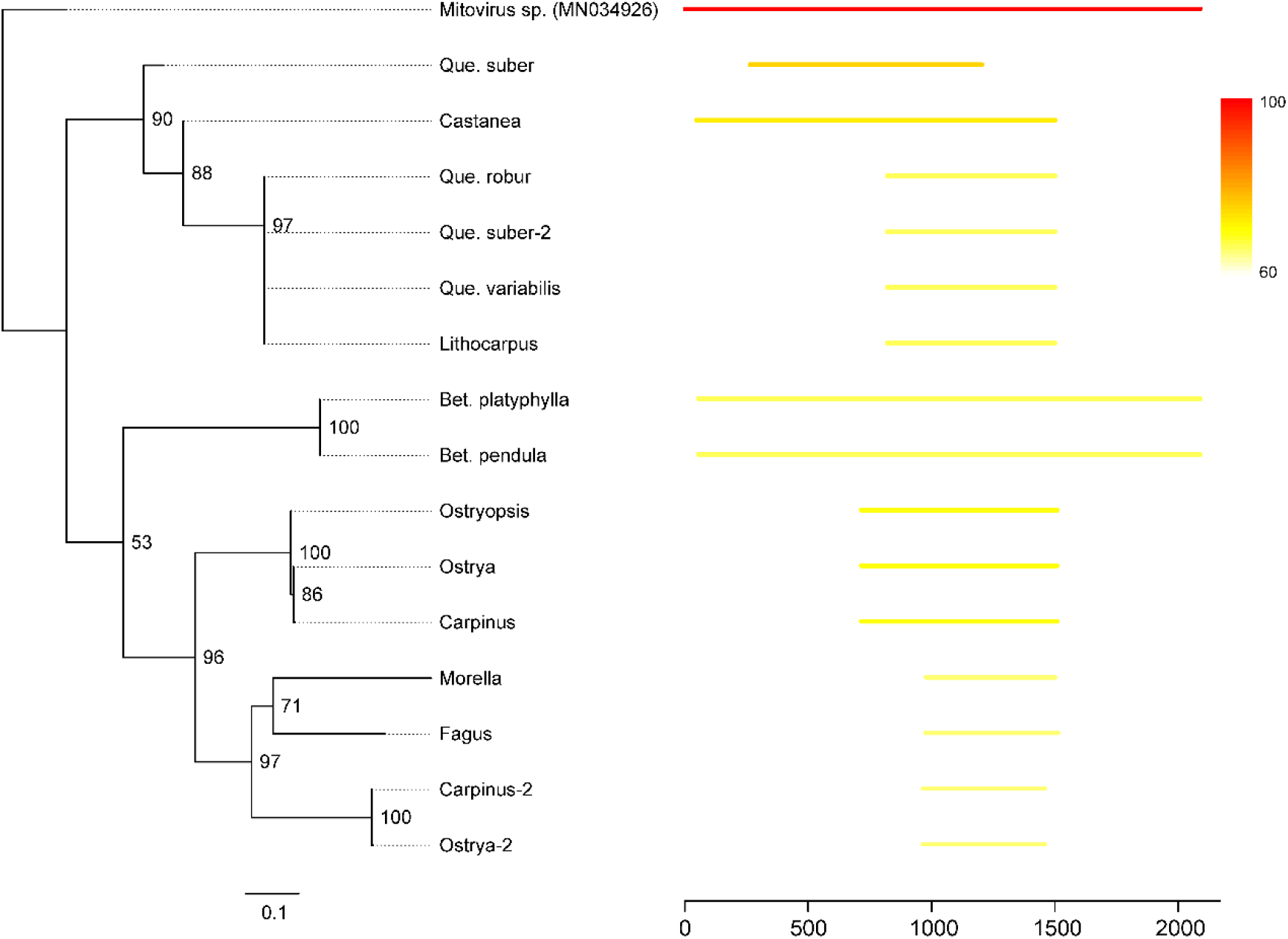
Mitovirus-like sequences in Fagales and their relationship. Lines on the right side shows the position and similarity of the hits against *Mitovirus* (MN034926). Only hits longer than 400 bp were shown. The tree on the left side was constructed by the hit sequences with ML method.

Fagales belong to the “nitrogen-fixing lineage”. At least three species in our sampling have the ability of nitrogen fixation: *Casuarina*, *Morella* and *Alnus* (Yelenik and D’Antonio, 2013; Huisman and Geurts, 2020). No evidence showed they contain sequences more similar to bacteria.

## Discussion

### A visual method helps precise assembly

In this study, we summarized the potential problems during mitogenome assembly and gave a credible strategy to obtain high-quality mitogenomes. Via visual processes in the powerful software Geneious, it can make sure every base correct (SNPs can be seen as well, if it has). That is necessary when we what to explore more detailed studies of DNA molecular evolution, such as the RNA editing, intron splicing and DNA double strand breaks and repairs. Our strategy is based on NGS short reads. A great benefit of short reads is the stock from other sequencing projects or on public databases (NCBI or CNGBdb).

An obvious disadvantage of short reads is they can’t overcome long repeats. So, the connection around long repeats of our assemblies is most likely pseudo or can only represent one potential type. A potential way to avoid mis-assemblies is checking the completeness of the gene colinear blocks (Chaw et al., 2008), like *nad1*e4-*matR*-*nad1*e5 in Fagales. Considering that plant mitogenomes rearrange more frequently as long repeats (Kozik et al., 2019), whether the genome has a "standard" structure is largely uncertain. The architecture of plant mitogenome is mysterious. Despite of the complex structure of mitogenome under microscopy (Backert and Börner, 2000; Manchekar et al., 2006; Cheng et al., 2017), we showed that most mitogenomes can still be assembled as circle(s).

Intracellular gene transfer (IGT) between genome compartments is a common phenomenon. Besides of MTPTs, plastid fragments can also transfer to the nucleus, referred to as nuclear plastid insertions (NUPTs, like the plasmid in *Carpinus*). Likewise, there are mitochondrial nuclear insertions (MTNUs) and nuclear mitochondrial insertions (NUMTs). Precise assembly of them should be the first step to solve the genome interaction and evolution. Unfortunately, with the increase of genomic studies and development of the sequencing technology, this issue is still neglected in almost all projects. Nuclear genomes always contaminated with true mitochondrial contigs (e.g., Alverson et al., 2011b). Therefore, interactions between nucleus and mitochondrion were not analyzed in this study. We hope our method of assembling MTPTs can give some hints to the others, and furthermore find better ideas and crack other IGTs.

### How mitogenome evolves?

Size variation between close species is a common feature for the plant mitogenomes, *e.g.*, *Viscum album* and *V. scurruloideum* (Petersen et al., 2015; Skippington et al., 2015), *Sliene conica* and *S. noctiflora* (Wu et al., 2015; Wu and Sloan, 2018), *Cucumis melo* and *C. sativus* (Alverson et al., 2011a; Rodríguez-Moreno et al., 2011). The reasons could be complex.

Duplications, IGT and foreign DNA all contribute to the expansion (Alverson et al., 2011a; Rice et al., 2013). In Fagales, we also saw size variation. Peculiarly, *Carpinus* is notable larger than relatives. However, its lengths of repeats, MTPTs as well as genus-specific sequences all failed to explain the size divergence. Only one possibility remains: it has more homologs with other Fagales. It was proved by searching the homologs between *Carpinus* and other lineages (Figure S5). That is actually quite confusing. What’s the ancestor of Fagales like? Did it have a big mitogenome, and lost different sequences independently in other species? The model was used to explain the size variation in kiwifruits, which also have a similar situation (Wang et al., 2019). But in Fagales, it’s too exaggerated if so many species had independent losses only without *Carpinus*. If no so, why only *Carpinus* has more homologs with other families? How the sequences spread?

Mitochondrial plasmids are small autonomous circular or linear extrachromosomal DNA in mitochondria. So far, it was found in many species, including maize, rice and carrot (McDermott et al., 2008). Their origin and function remain mysterious. The plasmid found in *Carpinus* gives a clear evidence that nuclear DNA is one of the sources. The GC content of plasmid if much lower than other mtDNA. Plasmid might be only an intermedium before the foreign DNA really compatible by the main chromosome. A fully understanding of this may need a better assembly of MTNUs and NUMTs.

Our genus-specific DNA analysis exhibit mitogenome in Fagales (or probably all angiosperms) is very mosaic. All the species have sequences that most similar to distant taxa in seed plants, including gymnosperms. We even don’t know if we should call them “HGT”, since it seems so common. Some orders have distinctly more similar sequences, and none of them really close to Fagales (Figure 4, *e.g.*, Amborellales, Lamiales and Ericales). *Amborella* has extreme HGTs from many species, including Fagales (Rice et al., 2013). We showed that these HGTs were mainly shared with Casuarinaceae. Since we used genus-specific DNA, the direction of these HGTs is actually uncertain. Parasitic plants are famous for massive HGTs from hosts (Bellot et al., 2016; Sanchez-Puerta et al., 2017; Kovar et al., 2018; Sanchez-Puerta et al., 2018). But in our analysis, parasitic plants didn’t have more HGT-like sequences with Fagales than other nonparasites (Table S4). The “wounding-HGT” model and “mitochondrial fusion occurs in a fundamentally similar manner” hypothesis were used to explain the HGT sequences between nonparasitic plants (Rice et al., 2013). It may also apply to Fagales cause no results were from other cellular organisms, even though some species are symbiosis with nitrogen-fixing bacteria. On the other hand, these HGT-like sequences in angiosperms may be just like the homologs we saw in *Carpinus*. Some DNA could spread in other unknown ways.

Mitovirus was only identified in fungi. But its sequence, especially RdRP region, are widespread in plant nuclear and mitochondrial genomes (Alverson et al., 2011a; Bruenn et al., 2015; Nibert, 2017; Silva et al., 2017; Chu et al., 2018; Nibert et al., 2018; Charon et al., 2020). The origin of plant mitovirus-like sequences was believed from plant pathogenic fungi through HGT (Bruenn et al., 2015). However, a direct HGT from fungi mitogenome is unlikely since the mitochondrial fusion between fungi and plants is difficult (Rice et al., 2013). What if the sequence transferred from fungi to nucleus first, subsequently from nucleus to mitochondrion? We searched the full-length mitovirus sequence of *Betula* mitogenome in *Bet. nana* and *Bet. pendula* nuclear genomes (Wang et al., 2013; Salojarvi et al., 2017), but only short pieces were found in *Bet. nana*. What’s more, the phylogeny of mitovirus-like sequences in Fagales doesn’t support they have the same origin (Figure 5; otherwise, the tree should have similar topology to plastid tree). We guess mitovirus can infect plants directly and frequently. Based on that, we infer the “third-part DNA”, including mitovirus and MTNUs, can account partially for the mosaic composition of plant mitogenomes. If two species get DNA from the same source, sometimes can make an illusion that similar sequences shared with far-away lineages; if different content was transferred in independent events, some species may share more homologs with others, like the situation in *Carpinus* (Figure 6). Since the transfers between the third-part and mitogenomes can happen independently and not limited to time, and mitogenomes themselves also encounter continuous rearrangements and deletion, time to time it will finally create extremely mosaic mitogenomes.

**Figure 6.**
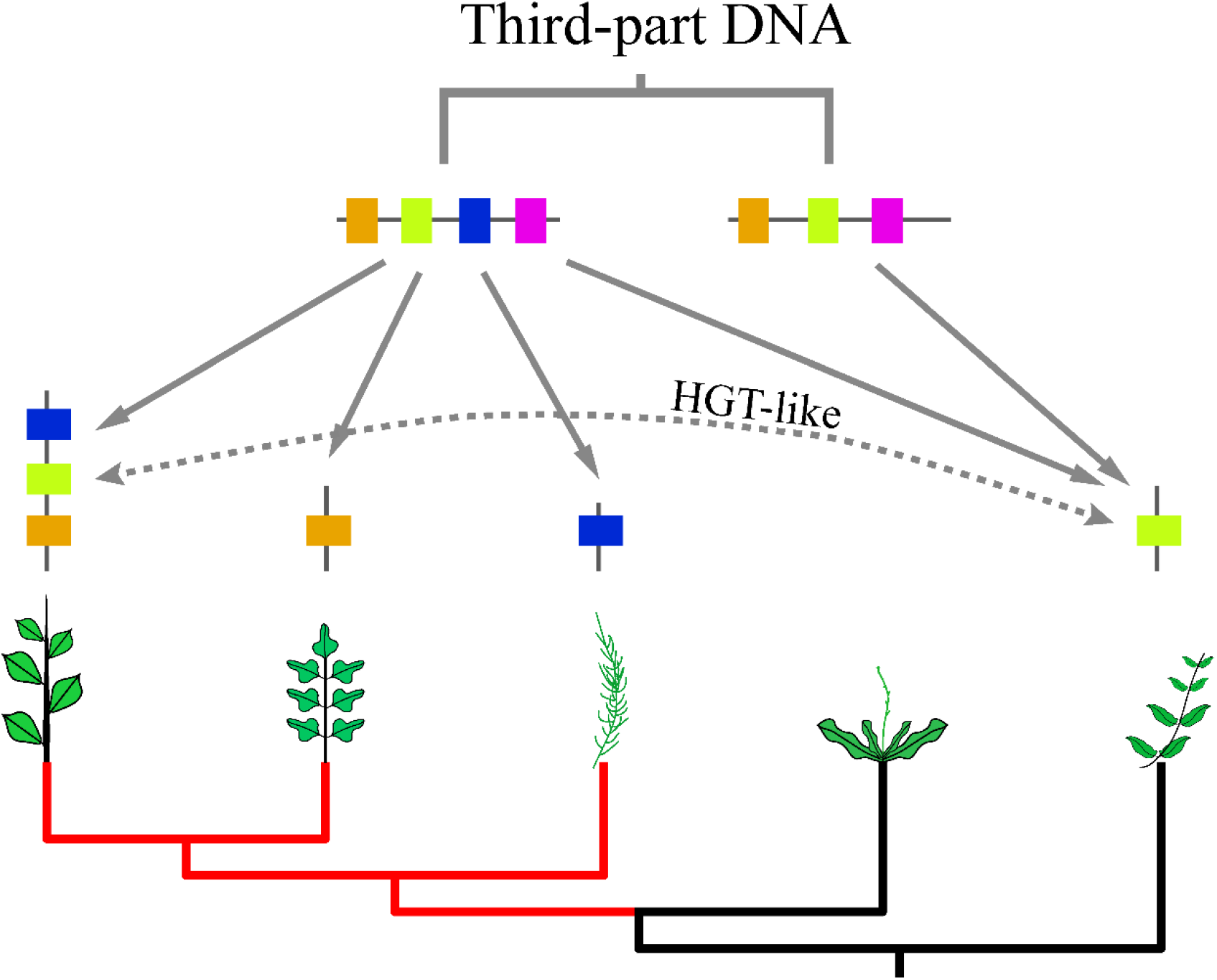
HGT-like sequences and inequal homologs caused by third-part DNA. Red branches show three close species. The colorful blocks indicate different genes or DNA fragments. The solid lines mean different transfer events. In the red lineage, the inequal transfers will make some species has the more homologs. The dot line means it creates an HGT-like sequence if one gene was transferred two independent times in distant lineages.

## Methods

### Species and Data acquisition

DNA sources of Fagales was retrieved from NCBI Sequence Read Archive (SRA) database (https://www.ncbi.nlm.nih.gov/sra). At last, 23 species from 16 genera and 5 families were chosen to do assembly (Table S1). All the data we used was whole genome sequencing (WGS), which means the reads contain genomes of all three compartments: nucleus, mitochondrion and plastid (including chloroplast). In plant cell, normally organelle genomes have much higher copy number but much smaller size than nuclear genome. It doesn’t need much data to get the mitogenome and plastome (Table S1).

### Genome assembly

After downloading and uncompressing, raw reads of each species were filtered low quality bases (PHRED < 30) by Trimmomatic v0.36 (Bolger et al., 2014). Clean reads *ca.* 2-4 Gb were used to perform *do novo* assembly via SPAdes v3.13 (Bankevich et al., 2012) (Table S1). To get the plastomes of *Casuarina equisetifolia*, *Lithocarpus fenestratus* and *Quercus suber*, plastid contigs of each was determined by BLASTN (Camacho et al., 2009) searching all assembled contigs against *Betula pendula* plastome (GenBank ID: NC_044852). Then we mapped clean reads to plastid contigs in Geneious R10 (Biomatters, Inc.), and extended and connected the contigs manually until they can join into one. IR boundaries were found by searching repeats via Geneious “Repeat Finder” plugin. Mitogenome is normally more variable in terms of content and structure. To figure out its contigs, we firstly extracted a preliminary set from total contigs by BLASTN using *Betula pendula* mitogenome (GenBank ID: LT855379) as reference (word size: 16, evalue: 1e-20), and extracted all hit sequences longer than 500 bp. Afterwards, we used two ways in Geneious to narrow it down. 1) Contigs were annotated via Geneious “Annotate from Database” function (here, the “Database” was comprised by all the known mitochondrial genes). If some gene was not found, reads were mapped to the gene to check the coverage, confirming its presence or absence. If the gene was missed from preliminary extraction, we would search the gene in all contigs and added the missing contigs. This way guaranteed the gene set of our assembly was complete. 2) We mapped the clean reads back to the chosen contigs in Geneious. Plastid (very high coverage) and other (unbalanced coverage) contigs were removed from the set. By that we got the approximate coverage of mitogenome. We used this coverage to bait other potential mitochondrial contigs from all contigs. The new chosen contigs were mapped back the reads again and removed non-mitochondrial contigs as before. This way guaranteed the DNA content of our assembly was (nearly) complete.

After got the mitochondrial contigs, next step was to connect them. Since connecting repeat ends is relatively easy, therefore we used a strategy solving them first, then connecting MTPT ends. Firstly, the repeats longer than 50 bp in the contigs were found through Geneious “Repeat Finder”. Then we mapped the paired reads to the contigs, and checked each repeat end and the corresponding pair. Repeat region can be seen from the coverage. Because of the high similarity, usually no MTPTs can be assembled directly in the contigs. We can only find them on plastomes (or plastid contigs). Unlike repeats, the coverage of plastome is commonly much higher than mitogenome, so it’s not easy to determine the MTPT regions by coverage. We fill the repeats and connected contigs at both sides. After that we mapped all the MTPT ends to the its plastome. The closest ends in right directions are most likely from the same MTPT. Sometimes rearrangement or recombination may also happen within MTPT, resulting a long distance or wrong direction (Figure S1f). In this case, paired reads can be used to find the right connection (Figure S1g-h). MTPTs and their plastid counterparts may not be 100% identical. Additional steps needed to correct the MTPTs we found from last step. Either we mapped reads again, checking and correcting the divergent bases manually (Figure S1c); or we used plastome filtered 100% identical reads, used the rest “Used reads” to mapped to mitogenomes again. The divergent base should be more clear (Figure S1d). Sometimes indels also exists (Figure S1e).

If it goes smoothly, after several map-check-connect iterations we can solve all the repetitive and MTPT ends, and get one or some circular chromosomes. At last, we mapped the PE reads again to check and correct the mis-assemblies - making sure every base is correct.

### Annotation

Annotations of the mitochondrial protein coding and rRNA genes were predicted by known mitochondrial genes by similarity in GENEIOUS, followed checking by eye and corrected manually. *Betula pendula* mitogenome was also annotated. tRNA were predicted using tRNAscan-SE v2.0 (Chan and Lowe, 2019). Coding genes with disrupted reading frames, premature stop codons, or non-triplet frameshifts were annotated as pseudogenes.

Total MTPT length was determined by BLASTN against a group of plastomes of Fagales (GenBank IDs were shown on Figure S4). Hits less than 100 bp were masked. Dispersed repeats within the genome were searched by BLASTN against itself. Hits with identity less than 95% were filtered. Repeat length was counted by custom Perl script. Only one part of the repeat pair was calculated and the overlapped bases were counted only once.

The annotations of the mitogenomes and their synteny were plotted by CIRCOS v0.69 (Krzywinski et al., 2009). Links were searched by BLASTN with default parameter and hit less than 500 bp were not considered. Homologs within the species were masked and only those between species were used.

### Phylogeny

The substitution rate of mitochondrial genes is very slow (Palmer and Herbon, 1988). But it contains hundreds RNA edit sites (Small et al., 2020). To prevent potential influence, now a typical way is removing these sites. Plastid RNA edit sites were ignored because of its small amount. All mitochondrial protein coding sequences (CDSs) were extract and aligned by mafft with “auto” mode. RNA edit sites were predicted by PREP website (Mower, 2009). If a codon has edit site(s) in one species, the codon of all the species were deleted. At last, the aligned genes were concatenated into one matrix and build the maximum likelihood (ML) tree on CIPRES v3.3 (Miller et al., 2010) with RAxML-HPC2 on XSEDE (Stamatakis, 2014), under the GTRGAMMA model and bootstrap iteration 1000. Plastid CDSs were aligned and build the tree in a same way just without removing RNA edit sites.

### Genus-specific DNA Analysis

Each mitogenome was blasted against a database made by all Fagales mitogenomes with evalue 1e-5 and word size 16. Then genus-specific sequences, *i.e.*, sequences only exist in this genus in Fagales, longer than 300 bp were isolated by custom Python script. Considering the non-monophyletic relationship of *Quercus* (Figure S4), *Que. suber* and *Que. variabilis* were seen as one genus excluding *Que. robur.* Afterwards, each genus-specific DNA was blasted against NCBI nr database online with the same parameter and saved the first 100 targets. Best-hit(s) of each genus-specific sequence were examined (more than one best-hits could exist in one sequence if they hit different area) by another Python script. Only the best-hits longer than 100 bp were counted. MTPTs were removed from the results. Subsequently, grouped these best-hits into orders and plotted in R. Ape package cophyloplot function were used to plot the face-to-face tree (Paradis et al., 2004) and the connections were colored by RColorBrewer (https://colorbrewer2.org/). The position of the orders was referred to the Angiosperm Phylogeny Group website (Stevens, 2001 onwards).

## Availability of supporting data

**Table S1.** Used SRA data.

**Table S2.** Mitogenomes with linear structure and potential issues of them

**Table S3.** The number of dispersed repeats and MTPTs in mitogenomes. Repeat number was grouped by length.

**Table S4.** Best-hits regions and potential original orders.

**Table S5.** Lengths of total genes-specific DNA, best-hits and potential origins.

**Figure S1:** Additional steps to get MTPT sequences. **A-E**: two methods to get exact MTPT bases. **C-D** show divergent bases between plastid and MTPT sequences, **E** shows two indels (when it exists, you can see obvious unmatched bases at two sides); **F-H**: a way solving nonadjacent MTPT ends. The direction of the arrow points to the direction the end should extend to. U1, U2: “unused reads” from Geneious custom sensitivity mapping.

**Figure S2:** A case to show the repeat coverage problem hinders a circular assembly. Red arrows indicate the repetitive end at the front and the position and direction of the other repeat pair. We can see higher coverage in the direction it should extend, but the coverage is gradually down to the normal level. Blue arrows are same but for the tail.

**Figure S3:** Circos plots of each family. **(a)** Betulaceae; **(b)** Casuarinaceae; **(c)** Fagaceae; **(d)** Juglandaceae. Links within each species were not plotted. The outer ring shows the position of protein genes and rRNA (red), tRNA (blue), repeats (yellow) and MTPTs (grey). It’s impossible to trace every link because of the density. But we can see the similarity between species and which has more species-specific sequences within each family.

**Figure S4:** Phylogenetic trees reconstructed by mitochondrial and plastid coding genes. **(a)**: mitochondrial tree; **(b)**: plastid tree.

**Figure S5.** Specific sequence length of *Carpinus* **(a)** and *Ostrya* **(b)** and other Fagales. The number in each circle means the specific sequence length of *Carpinus* **(a)** or *Ostrya* **(b)** and all the species within the circle. For example, the Casuarinaceae circle in **(a)** indicates *Carpinus* has 356 Kb sequences homologous only with any species of Casuarinaceae and other Betulaceae, but not with other species out of the circle. The comparison between **(a)** and **(b)** reveals the big size of *Carpinus* is mainly attributed to more homologous DNA with other Fagales.

## Data and Code Availability

The assembled genomes have been deposited to CNGBdb under Project CNP0001491 (mitogenomes: accessions N_000011064 - N_000011115; plastomes: accessions N_000011061 - N_000011063; Carpinus mitochondrial plasmid: accession N_000011116). The used scripts can be found in Github (https://github.com/fengyanlei33/Fagales_mitogenome).

## Acknowledgements

We gratefully acknowledge Xingxing Shen (Zhejiang University) for comments and suggestions.

