## Supplementary figures and images for "An updated assembly strategy helps parsing the cryptic mitochondrial genome evolution in plants"

### Supplemental Figure 1

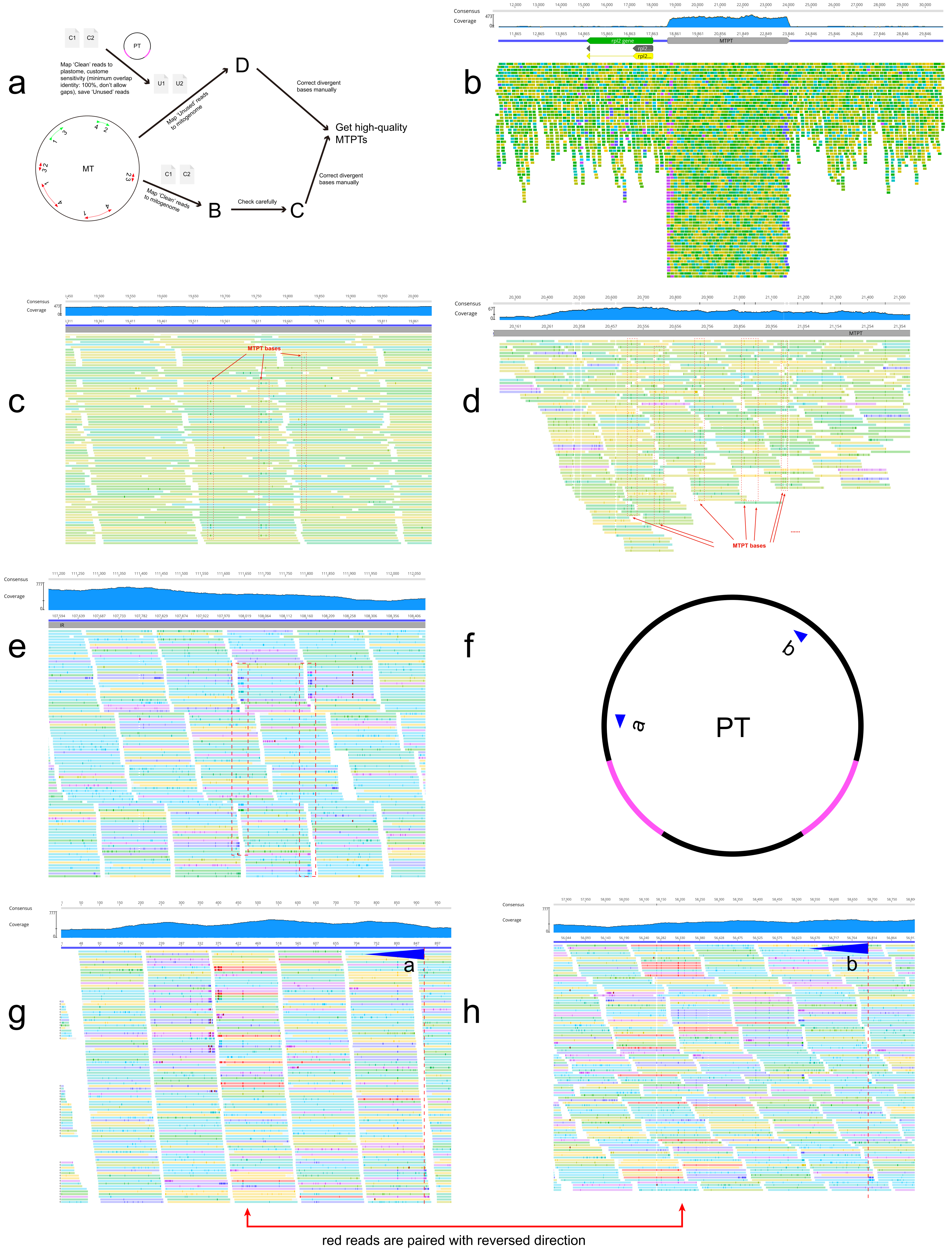

### Supplemental Figure 2

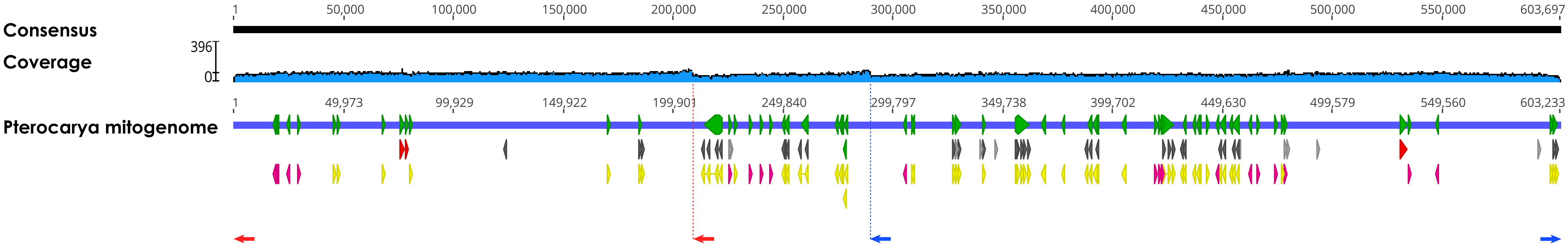

### Supplemental Figure 3

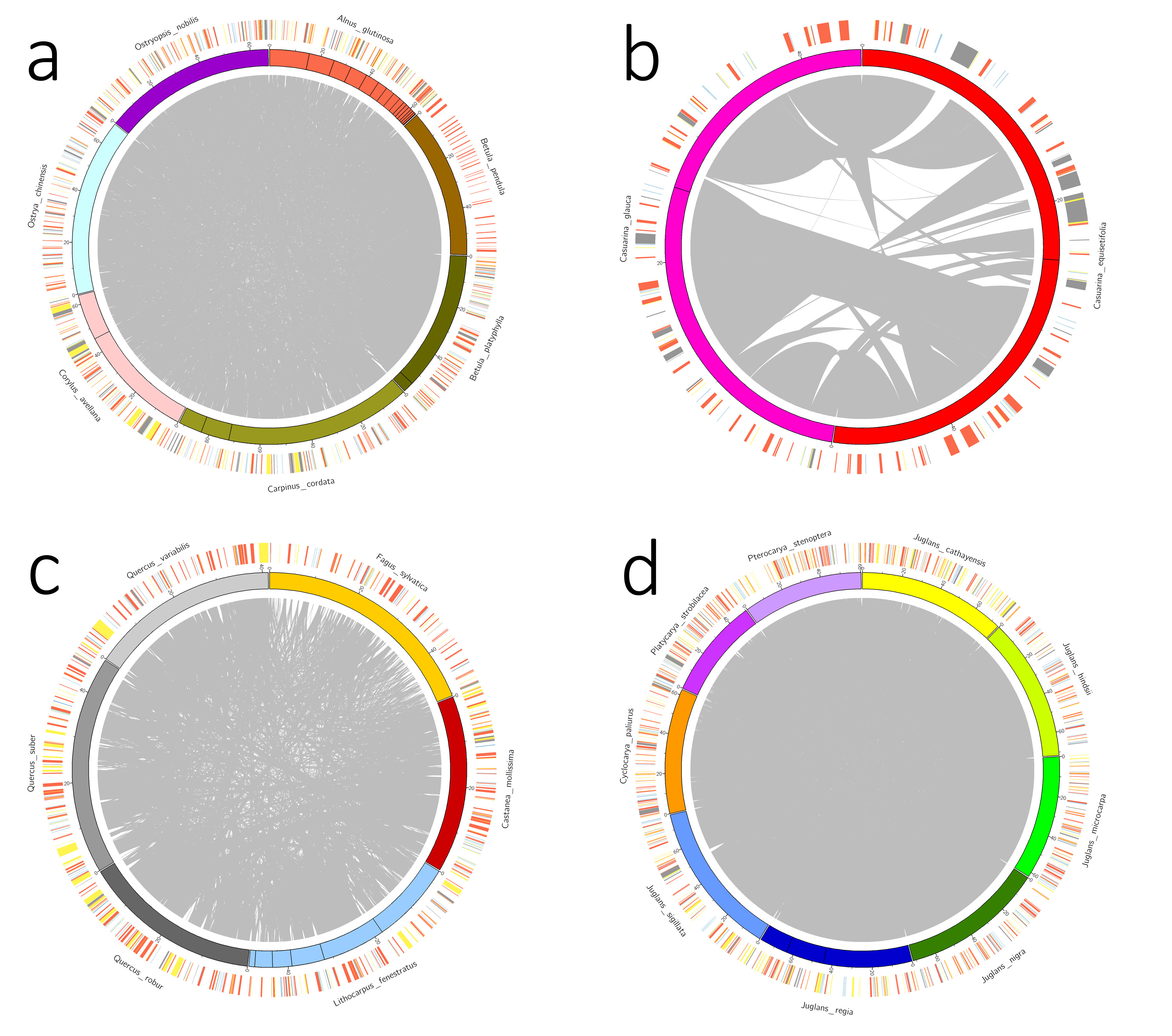

### Supplemental Figure 4

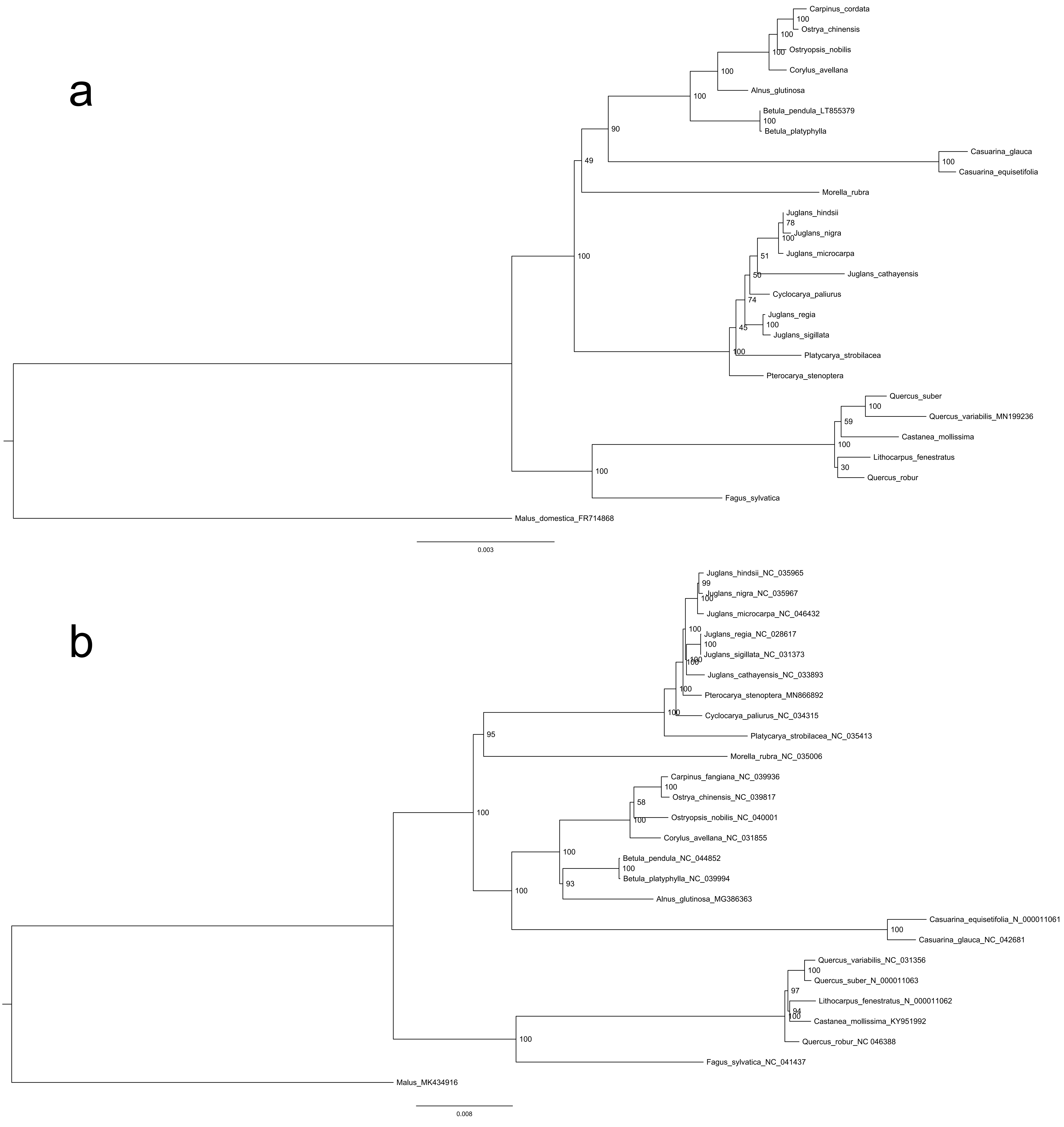

### Supplemental Figure 5

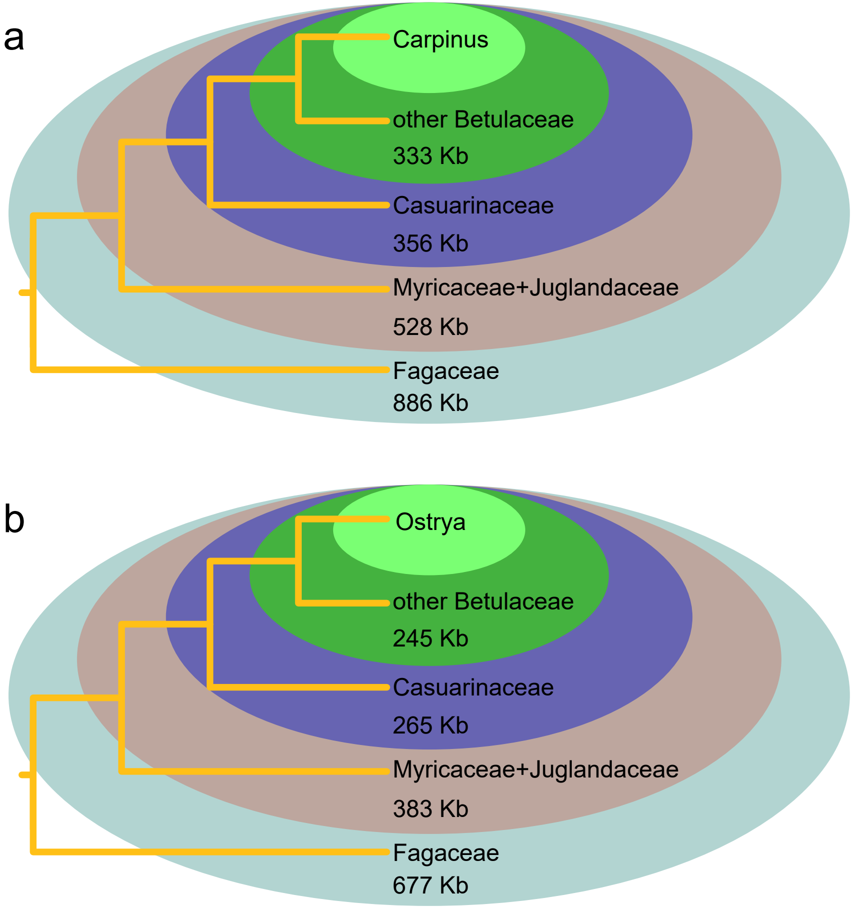
